# Characterizing multiple dimensions of climate hazards for conservation planning: a case study of estuaries in the Pacific Northwest

**DOI:** 10.64898/2026.08.03.741880

**Authors:** Rekha Marcus, Nancy Shackelford, Gerald Singh

## Abstract

Incorporating the complexities of climate change into conservation planning can be challenging. Climate change is projected to change the mean and variability of temperature and precipitation, and change dynamics of extreme events of many climate variables globally. These changes will all have compounding effects on ecosystems worldwide, affecting species distributions, seasonal timings, and population dynamics. However, most recent climate-informed conservation planning frameworks only focus on changes in mean conditions, leaving out ecologically important information about seasonality and extreme events. Using a case study of estuaries in the Pacific Northwest, this research seeks to answer the question “How can the effects of climate change on estuaries best be modelled and described for practical use in ecological management?” To answer this question, we used downscaled climate projection data to estimate changes in the mean and variability of temperature and precipitation. Applying extreme value theory to these projection data, we also projected changes in the magnitude of extremes in temperature and precipitation for estuaries in the region. Using descriptive statistics and open source data, our results present a novel, holistic method of understanding climate risk, including estimating extreme events, highlighting a key research gap in conservation planning. These results also highlight that trends in the mean, variability, and magnitude of climate extremes are not consistent with each other, further underscoring the importance of considering multiple dimensions of climate change together. Despite uncertainty given by climate models, the methods presented here provide a reasonable approach to plan for conservation management in the face of climate uncertainty.

## Introduction

The complex nature of climate change makes it difficult to incorporate into conservation planning (Belote et al., 2017, 2018; Chung et al., 2025; Elsen et al., 2020; Marcus & Noonan, 2023). The term “climate change” often encompasses consideration of changes in the abiotic environment and generation of hazards such as changes in mean climate characteristics, seasonal variability, and increases in both frequency and intensity of extreme events around the world (Calvin et al., 2023; Field et al., 2012). Oftentimes, conservation planners must worry about immediate issues such as loss of funding or project relevancy, making complex, long-term challenges like climate change lower priorities for decision-making (Jewell et al., 2023). However, there is increased recognition that ignoring climate impacts can render conservation plans obsolete or ineffective (Groves et al., 2012; Pressey et al., 2007; Reside et al., 2018; Stralberg et al., 2026).

Many climate-inclusive conservation planning strategies focus primarily on adapting and planning for changes in mean conditions (Hannah et al., 2002a; Hansen et al., 2010; Thompson et al., 2013). However, changes in mean climate conditions don’t capture the full range—or even the most important—of climate effects on the environment (Merder et al., 2026; Thompson et al., 2013). Changes in the climate system will lead to changes in the shape of the distribution of many different climate variables, leading to changes in seasonal timings and the frequency, intensity, and timing of extreme weather events (Caparros-Santiago et al., 2021; Diffenbaugh et al., 2017; Field et al., 2012; Ghil & Lucarini, 2020; Pendergrass et al., 2017; Stouffer & Wetherald, 2007; Visser & Both, 2005). These changes are expected to be biologically more significant than mean changes, as many species rely on seasonal patterns for food availability, and extreme events can have significant effects on biodiversity (Fusi et al., 2022; García-Carreras & Reuman, 2013; Menzel et al., 2006; Visser & Both, 2005). While recent years have brought to the public attention the significance of considering variability and extreme events in climate change, these are yet underrepresented in conservation management.

Seasonal changes and environmental variability drive habitat selection, migration patterns, and reproductive strategies for most species (Caparros-Santiago et al., 2021; Dagtekin et al., 2024; de Zwaan et al., 2022; Johns et al., 2025; Visser, 2016; Williams & Middleton, 2008), making climate variability an important consideration for conservation planning. Ecologically, changes in variability of specific climate variables such as temperature are linked to changes in seasonal timing, which are significant for ecosystem functioning. For example, early spring events change the spatial variability of plants, leading to mismatches in the distributions of species and their food (Menzel et al., 2006; Visser & Both, 2005). At the ecosystem level, environmental variability is linked to resource availability (eg. Baert et al., 2022; Clulow et al., 2011), and population growth and decline (Easterling et al., 2000; Williams & Middleton, 2008). As climate change drives changes in seasonal timings and increases the variability of important climate variables (Stouffer & Wetherald, 2007; Thornton et al., 2014; Visser & Both, 2005), this necessitates understanding changes in the shape of the distribution of these variables.

Beyond changes in seasonality, preparedness for extreme events is being recognized as an important facet of conservation planning (K. R. Jones et al., 2016; Ly & Diffenbaugh, 2025; Maxwell et al., 2019). Extreme events are projected to increase in both frequency and magnitude globally (Diffenbaugh et al., 2017; Field et al., 2012), including extreme temperatures, and increased storm behaviour. As they are rare by nature, extreme events can be difficult to model and plan for, but are equally as ecologically significant as changes in variability. Extreme heat waves have been shown to result in species mortality across many taxonomic levels (Jones, 2018), disrupting food webs and causing extreme stress to organisms (Anadón et al., 2024; L. A. Jones et al., 2020; Pardo et al., 2017; Pérez-Romero et al., 2019; Piatt et al., 2020; Shields et al., 2019; Williams & Middleton, 2008). Extreme rainfall can lead to flooding, especially in areas where natural flooding regimes have been restricted (*e.g.*, dams or channelized streams) (Reich & Lake, 2015), which in turn can destroy healthy habitat, introducing toxins into the watershed and exacerbating the already shrinking habitat for many species (Jentsch & Beierkuhnlein, 2008; Liu et al., 2021).

In the northern Pacific coast of North America (here termed Pacific Northwest), an ecosystem that is a priority for many conservation planners is estuaries. These estuaries are uniquely variable ecosystems that occur on coasts all around the world, and are generally defined by a mix of freshwater input from a river or runoff, and saltwater input from an adjacent marine ecosystem (Pritchard, 1967). These ecosystems provide habitat for 80 percent of coastal species in at least one life stage; this includes bioculturally significant species such as southern resident killer whales (*Orcinus orca*), anadromous salmon (*Oncorhynchus spp.*), and billions of migratory birds (Kelsey, 1995). By nature, estuaries are open ecosystems that rely completely on the existence of other ecosystems, and given their connection to tidal regimes, they are highly variable and thus necessitate species that are adapted to variability (McLusky & Elliott, 2004; Whitfield & Elliott, 2011). However, as climate change drives fast regime shifts in the climate conditions of these estuaries, the ecosystems themselves and the species that inhabit them may not be able to adapt quickly enough to avoid negative impacts. Thus, regional estuary conservation planners could use detailed, local-scale information on climate impacts in order to make the best decisions for these ecosystems. The purpose of this research is to address this important gap. This research seeks to answer the question “How can the effects of climate change on estuaries best be modelled and described for practical use in ecological management?”

To answer the research question, using estuaries on the Pacific Northwest coast as a case study to examine changes in climate conditions, this work will use existing projection data for multiple future climate scenarios to estimate for each estuary the change in (a) mean temperature and precipitation (b) variability in temperature and precipitation and (c) magnitude of extreme events in temperature and precipitation. Current global models predict an increase in both mean temperature and mean precipitation for the Pacific Northwest (Calvin et al., 2023; Intergovernmental Panel On Climate Change (IPCC), 2023). However, knowledge on temperature and precipitation variability and extremes is limited for estuaries in this region (eg. Bashevkin & Mahardja, 2022; Gross et al., 2023; Jarrin et al., 2022; Shellenbarger & Schoellhamer, 2011). Because estuaries in the Pacific Northwest provide home for so many migratory species, incorporating this knowledge into regional scale conservation planning efforts to help ensure adequate habitat is maintained along this migratory pathway is essential, especially as species shift their ranges to adapt to climate change (Hannah & Midgley, 2023).

## Methods

### Study area

The study area, referred to here as the Pacific Northwest, covers the coastal portion of western North America coincident with the Marine West Coast Forests ecoregion (Commission for Environmental Cooperation (CEC), 1997). The study area covers 344,023,946 square kilometres from the northern coast of California, and the entire Pacific coast of Oregon, Washington, and British Columbia (between bounding box coordinates -133.13 and -121.93 [longitude] and 38.75 and 55.93 [latitude]; Figure 1), representing almost the entirety of the ecoregion where estuaries have been described and mapped in detail (Pacific Birds Habitat Joint Venture Technical Team, 2019; Pacific Marine and Estuarine Fish Habitat Partnership, 2018).

**Figure 1:**
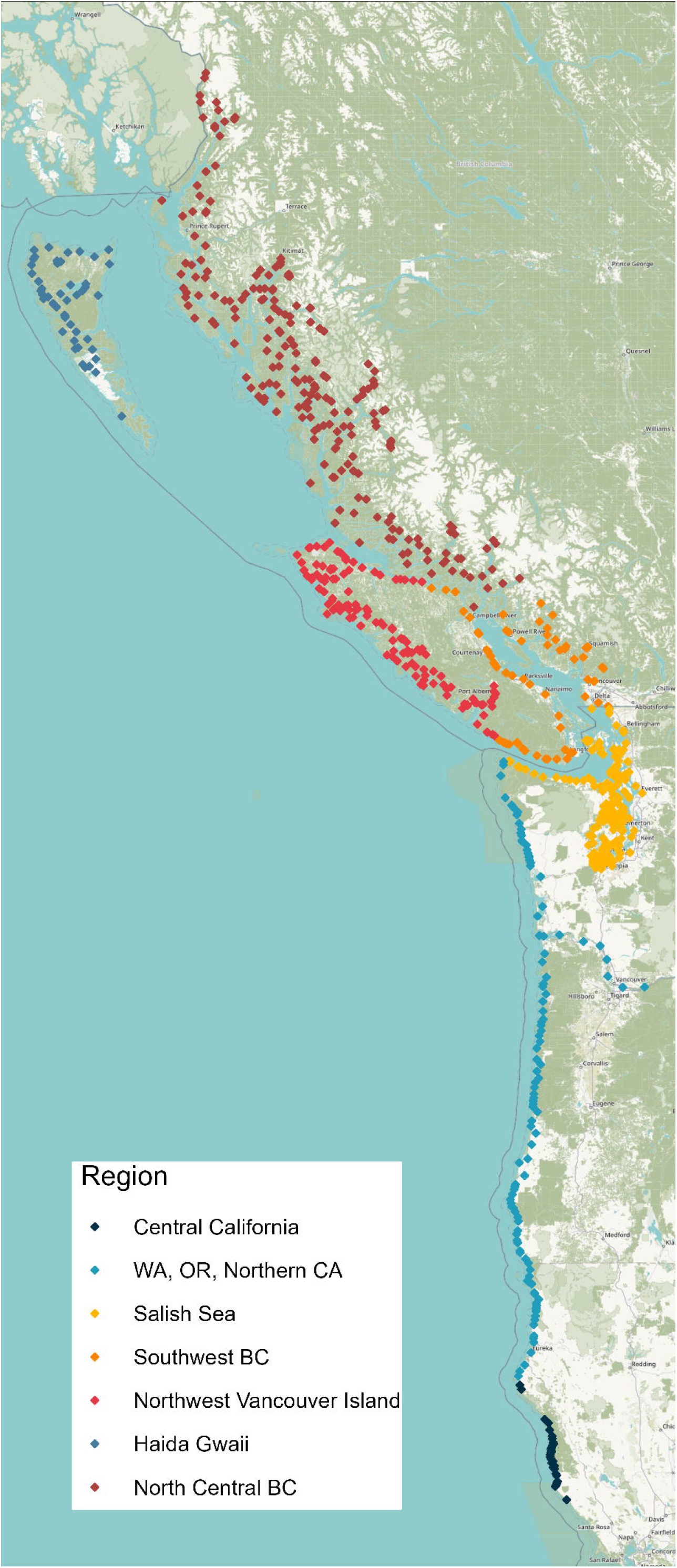
Map highlighting all estuaries within the study region (n = 749). Points denote estuaries, and colors correspond to the ecoregion the estuary belongs to. Map lines delineate study areas and do not necessarily depict accepted national boundaries.

The study area contains 749 estuaries, ranging in size from less than 1 km^2^ to 660,000 km^2 .^(Pacific Birds Habitat Joint Venture Technical Team, 2019; Pacific Marine and Estuarine Fish Habitat Partnership, 2018). These estuaries can be divided into subregions, which correspond to unique ecoregions found in the Pacific Northwest ecozone from north to south: North Central British Columbia; Haida Gwaii; Northwest Vancouver Island; Southwest British Columbia; Salish Sea, Washington; coastal Washington, Oregon, and northern California; and Central California (Demarchi, n.d.; Griffith et al., 2016; Pater et al., n.d., Figure 1).

### Data

In order to answer the research question, we first determined which climate variables would be most important to understand at the regional scale. We then collected both historical and future projection data for the identified variables, cropping them to each estuary in the study area (n = 749).

Changes in temperature that accompany climate change will affect many of the qualities of estuaries that make them valuable habitats. As these are often ecosystems of shallow water, they are more susceptible to warming, and they are warming at a rate faster than adjacent ocean or riparian ecosystems (Scanes et al., 2020). These changes in temperature can affect primary productivity and growth of plants essential to these ecosystems, as well as lead to range shifts in communities of different organisms (Lonhart et al., 2019). This can, in turn, affect recruitment, community composition, and the ability of these ecosystems to act as nurseries for many species that rely on them during breeding (Cloern et al., 2016; van Niekerk et al., 2022). Along with changes in temperature, climate change will affect major precipitation patterns. Precipitation is linked to salinity patterns in estuaries, which is a major driver of community composition (Chilton et al., 2021; Cloern et al., 2016). Additionally, increases or decreases in precipitation will drive parallel changes in streamflow, which will drive further habitat loss, changes in water quality, sedimentation patterns, and primary productivity, and will change migration patterns of organisms such as migratory birds and anadromous salmon (Chilton et al., 2021). Temperature and precipitation both can be observed at the global scale, but also generate significant effects at the local scale. Because changes in temperature and precipitation will affect estuaries in various ways, they were chosen as the two variables to include in this analysis.

Downscaled climate data used in this analysis were extracted from the Climatologies at a High resolution for Earth’s Land Surface Areas (CHELSA) project, which contains global historical and projected climatological data from world meteorological stations between 1979 and 2019, standardised and modelled at a ∼1km^2^ resolution, and is hosted by the Swiss Federal Institute for Forest, Snow and Landscape Research WSL (Karger et al., 2017). Our research used monthly historical means in temperature and monthly precipitation amounts from CHELSA, which are the most readily available and commonly used climate variables, and are biologically significant to estuarine productivity and functioning (Chilton et al., 2021; Emmett et al., 2000; Parker et al., 2019; Robins et al., 2016; Roessig et al., 2004). Future projection data for these variables used two Shared Socio-economic Pathway (SSP) scenarios, SSP1-2.6 and SSP5-8.5, which were chosen to represent the range of effects that climate systems may have on estuaries in the future. For each of these scenarios, data was acquired from 5 different climate projection models— GFDL-ESM4 (National Oceanic and Atmospheric Administration, USA), UKESM0-1-LL (Met Office, UK), MPI-ESM1-2-hr (Max Planck Institute, Germany), IPSL-CM6A-LR (Institut Pierre Simon Laplace, France), MRI-ESM2-0 (Meteorological Research Institute, Japan)—which was compiled and downscaled by CHELSA. Multi-model ensembles are commonly used in climate science in order to capture a wider range of potential futures, and avoid basing results on the specific assumptions made from a single model. By using climate projections from 5 different models for 2 different future scenarios, we hoped to better capture the variability in effects that estuaries may experience in order to better inform conservation decisions (see S3). Despite some criticism for the implausibility of SSP5-8.5 based on energy use and carbon emissions (and therefore the causes and radiative forcing; Pielke Jr et al., 2022), here we emphasize the effects of climate change, for which RCP 8.5 may serve as a worse case scenario (Pedersen et al., 2020). Table 1 details the layers extracted from this dataset for this project. Each of these data layers were cropped to the individual estuary level.

**Table 1:** Climate data layers used for this project. *These observational data were used for model validation only, as they are available only at a 0.5 degree resolution.

| Data Layer | Units | Spatial resolution | Model Source | Time Period |
| --- | --- | --- | --- | --- |
| Global Monthly Temperature | K*10 | 0.01° | CHELSA V2.1 | 1979 - 2019 |
| Global Monthly Precipitation | kg*100 | 0.01° | CHELSA V2.1 | 1979 - 2019 |
| Global Gridded Land Temperature (Monthly)* | K | 0.5° | NOAA<br>GHCN_CAMS | 1948 - present |
| Global Monthly Terrestrial Precipitation* | cm | 0.5° | NOAA University of Delaware | 1900 - 2017 |
| 30-year Mean Temperature | K | 0.01° | CHELSA CMIP6** | 2010 - 2100 |
| 30-year Mean Precipitation Amount | kg | 0.01° | CHELSA CMIP6 ** | 2010 - 2100 |
| 30-year Temperature Seasonality | Coefficient of Variation | 0.01° | CHELSA CMIP6 ** | 2010 - 2100 |
| 30-year Precipitation | Standard | 0.01° | CHELSA CMIP6 ** | 2010 - 2100 |
| Seasonality | Deviation |  |  |  |

Shapefiles for cropping estuaries in British Columbia were provided by the PBHJV, as part of their previous estuary mapping projects (Pacific Birds Habitat Joint Venture Technical Team, 2019). Shapefiles for cropping estuaries in the United States of America were obtained from the Pacific Marine and Estuarine Fish Habitat Partnership (PMEP)’s West Coast USA Current and Historical Estuary Extent dataset (PMEP, 2018).

### Statistical Analysis

In order to answer the research questions, we explored 3 parameters in the data: changes in mean, changes in variability, and changes in the magnitude of extreme events. These describe the changes in the distributions of climate variables that we expect to see under climate change (Field et al., 2012).

To measure relative changes in mean, we compared the historical monthly means of temperature and precipitation to the two future projected scenarios described previously. We calculated the historical baseline mean using available CHELSA monthly historical data from 1979 to 2019, and calculated the percent difference between this historical baseline and a projected 30-year mean (Figure 2a). Similarly, for changes in variability, we calculated the baseline historical variability as the standard deviation of the monthly means in the historic data between 1979 and 2019, and compared that to future 30-year seasonality (standard deviation) projections to find the percent change in variability (Figure 2b).

**Figure 2:**
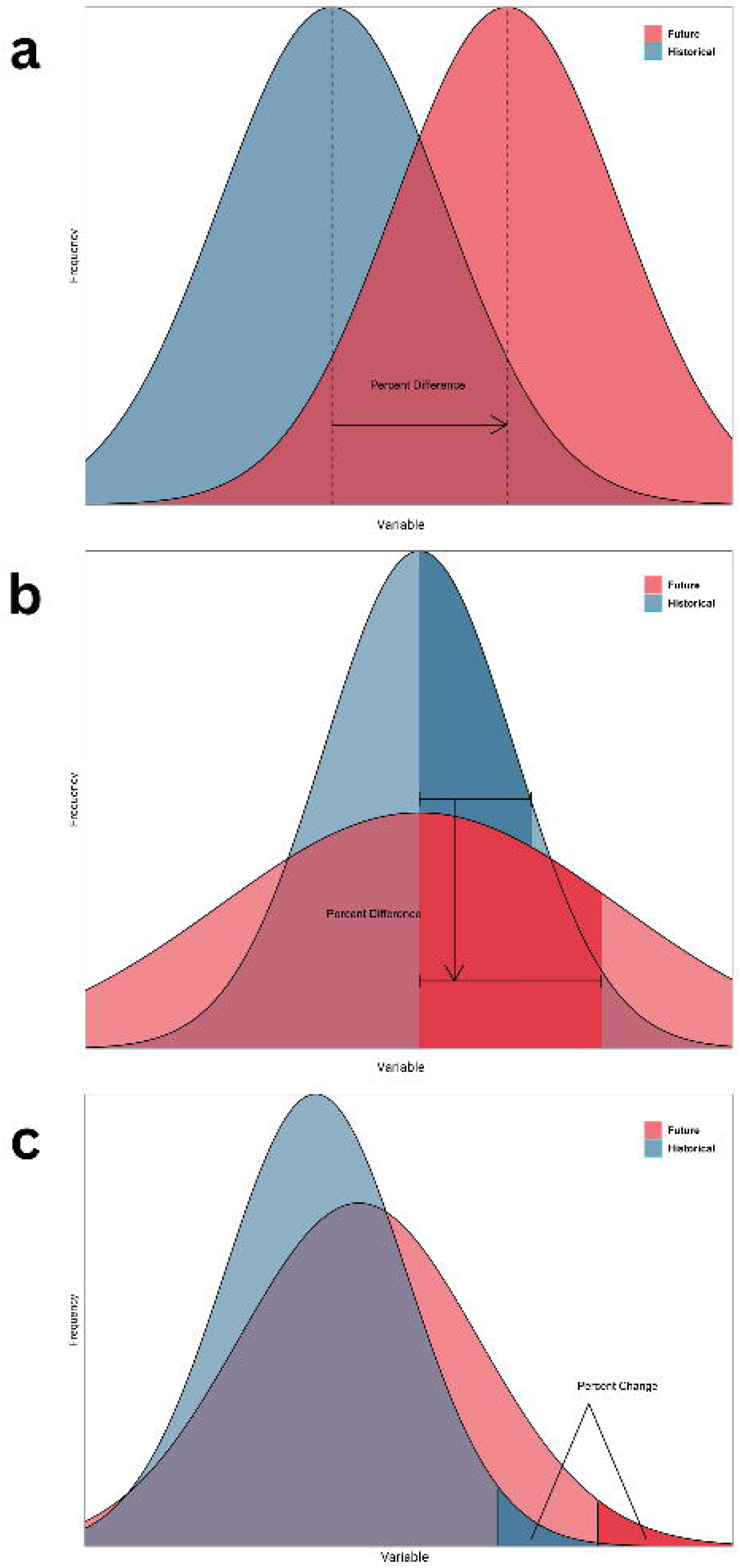
Conceptual figure demonstrating (a) the percent difference calculated between a historical and projected means (b) the percent difference calculated between historical and projected standard deviation, and (c) the percent difference between historical and projected expected shortfall, or area under the tails.

To estimate changes in the magnitude of extreme events, we used extreme value theory (EVT) to model behaviour of the data at the tails of their distributions. EVT fits the maximum or minimum values in a dataset to an extreme value distribution, which can then be used to describe trends in extreme values (Figure 3a, Fisher & Tippett, 1928). EVT is often used in climate research to approximate values such as the return period, or 1-in-n year event, estimating the magnitude of such an event by calculating a quantile value (Pausader et al., 2012). However, because the return period relies on a single value to capture the potential magnitude of an extreme event, financial analysis often makes use of another similar parameter, expected shortfall (ES), which measures the area under the tail and provides a more stable approximation of the risk of a given hazard (Hodge, 2000; Mak & Meng, 2014). While underapplied in climate and ecological methods (though see eg. Singh et al., 2024), ES could provide a more stable approximation of the effects of extreme temperature and precipitation events, mitigating some of the lack of robustness that exists in estimation of extremes.

**Figure 3:**
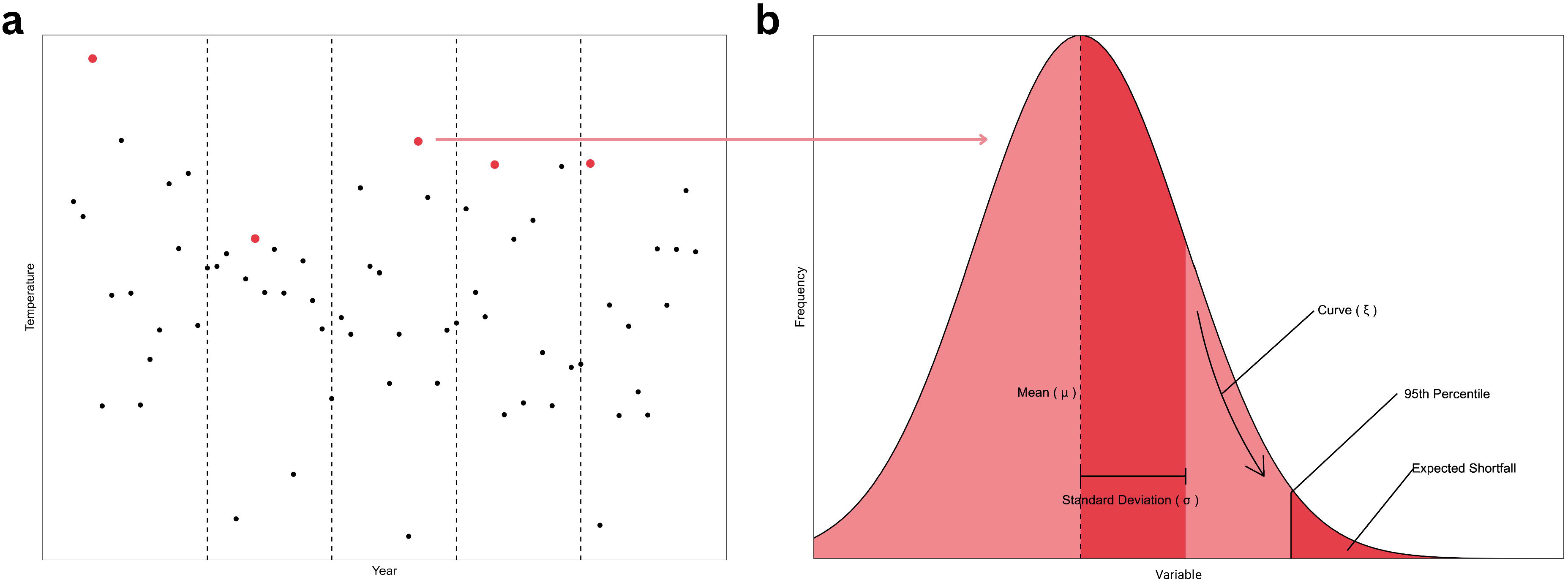
Conceptual figure demonstrating the construction and parameters of an extreme value distribution using the block maxima approach. Panel (a) highlights the maximum value in each block, which are then used to construct the distribution in (b), where the parameters of an extreme value distribution are displayed.

In order to calculate the ES of temperature and precipitation, we used the fevd function from the extRemes package in R (Gilleland, 2004) to fit a Weibull generalized extreme value distribution (EVD) to yearly block maxima (highest recorded) values in historical temperature and precipitation for each estuary (Fisher & Tippett, 1928, Figure 3a). The block maxima approach allowed for each estuary’s unique climate maxima to be captured, rather than relying on a single significant threshold that may or may not be ecologically relevant for estuaries across such a large latitudinal gradient, as would be required for the peaks-over-threshold approach to EVD. The EVD produced three parameters—location (μ, a measure of central tendency), scale (σ, a measure of spread), and shape of the tail (ξ) (Figure 3b) — to describe the behaviour of the historical temperature and precipitation data (Fisher & Tippett, 1928).

We then constructed an EVD for each modelled climate future (SSP 1 and SSP 5), using μ and σ values from projected future data, and ξ of the historical EVD. A critical assumption underlying this analysis is that the ξ parameter of the distribution would not change in the future. Because ES is less sensitive to changes in the tail shape than the return period (Fisher & Tippett, 1928; Hodge, 2000), we believe this assumption will still allow us to obtain a reasonable projection of future extreme events. To get an estimate of the magnitude of extreme events in both temperature and rainfall, we calculated the ES of both variables above the 95th and 99th percentile (extreme highs), using the qevd and pevd function from the extRemes package. The focus on extreme highs was chosen because both temperature and precipitation are projected to increase for the region, which we assume will lead to increases in the magnitude of extremes (Field et al., 2012; Masson-Delmotte et al., 2021), and extreme lows, especially of precipitation, are not as well captured by mean monthly data. Finally, for each estuary, we calculated the percent change between historical and future projected ES to determine how the magnitude of extreme events will change in different climate scenarios (Figure 2c). All analyses were conducted in R (v 4.4.3).

### Validation

The methods used to answer the research question relied on the use of modelled summary statistics (mean, standard deviation, and expected shortfall) to calculate changes in climate variables for estuaries. Because our approach relied on readily available summary statistics of modelled climate data rather than observed monthly or daily data points, a necessary piece of the method was validation of our approach. This is particularly important in our calculation of changes in the magnitude of extreme events, as we modelled these data ourselves using a novel method using summary statistics of mean and standard deviation. In order to understand how the use of both summary statistics and modelled data impacted the results, we calculated expected shortfall using 4 different datasets (described in Table 1):

1. Historical NOAA global climate data at a 0.5 degree resolution, which contained yearly observed precipitation and temperature data between 1908 and 2020;
2. Summary statistics (mean and standard deviation) of historical NOAA observed global climate data;
3. Historical CHELSA modelled global climate data at a 0.01 degree resolution, which contained monthly precipitation and temperature data between 1980 and 2021;
4. Summary statistics (mean and standard deviation) of historical CHELSA modelled global climate data;

Where the 1st dataset represents the “true” expected shortfall values, the 4th dataset includes the data used for generating our results for expected shortfall, and the 2nd and 3rd dataset provide a comparison to better understand the results of using modelled data and summary statistics. By using our described method to calculate the expected shortfall using modelled and raw data, both as summary statistics and as individual data points, we sought to explore whether the summarized modelled data could still capture the true value of expected shortfall for each estuary, and thus adequately approximate the effects of extreme climate events on estuaries.

## Results

### Model Validation

Figure 4 showcases the results of the model validation, done by calculating expected shortfall of both temperature and precipitation using different datasets. For temperature, there was a difference between the calculated extreme temperatures over the same time period when calculated based on observed data instead of modelled data; the observed data occasionally resulted in lower predicted extreme temperatures, although there was still significant overlap and the majority of the values calculated using modelled data were within one standard deviation of the observed data (Figure 4a). There was little difference in the distribution of residuals in the raw data versus the summary statistics data (mean raw value residuals = 1.69 ± 1.78, mean summary statistic residuals = 1.76 ± 1.83; Figure 4c); however, both distributions were centered around 1, indicating a slight overprediction of the magnitude of extreme temperature events based on our model (Figure 4c). To test for a significant difference between both groups, a Kruskal-Wallis test was calculated using a random subset of 30 estuaries and bootstrapped over 100 iterations (see S4 for more information), revealing a significant difference in the values calculated for expected shortfall of temperature across 4 different datasets (mean Kruskal-Wallis value (df = 3 , N = 4) = 23.38876, mean p-value < 0.001).

**Figure 4:**
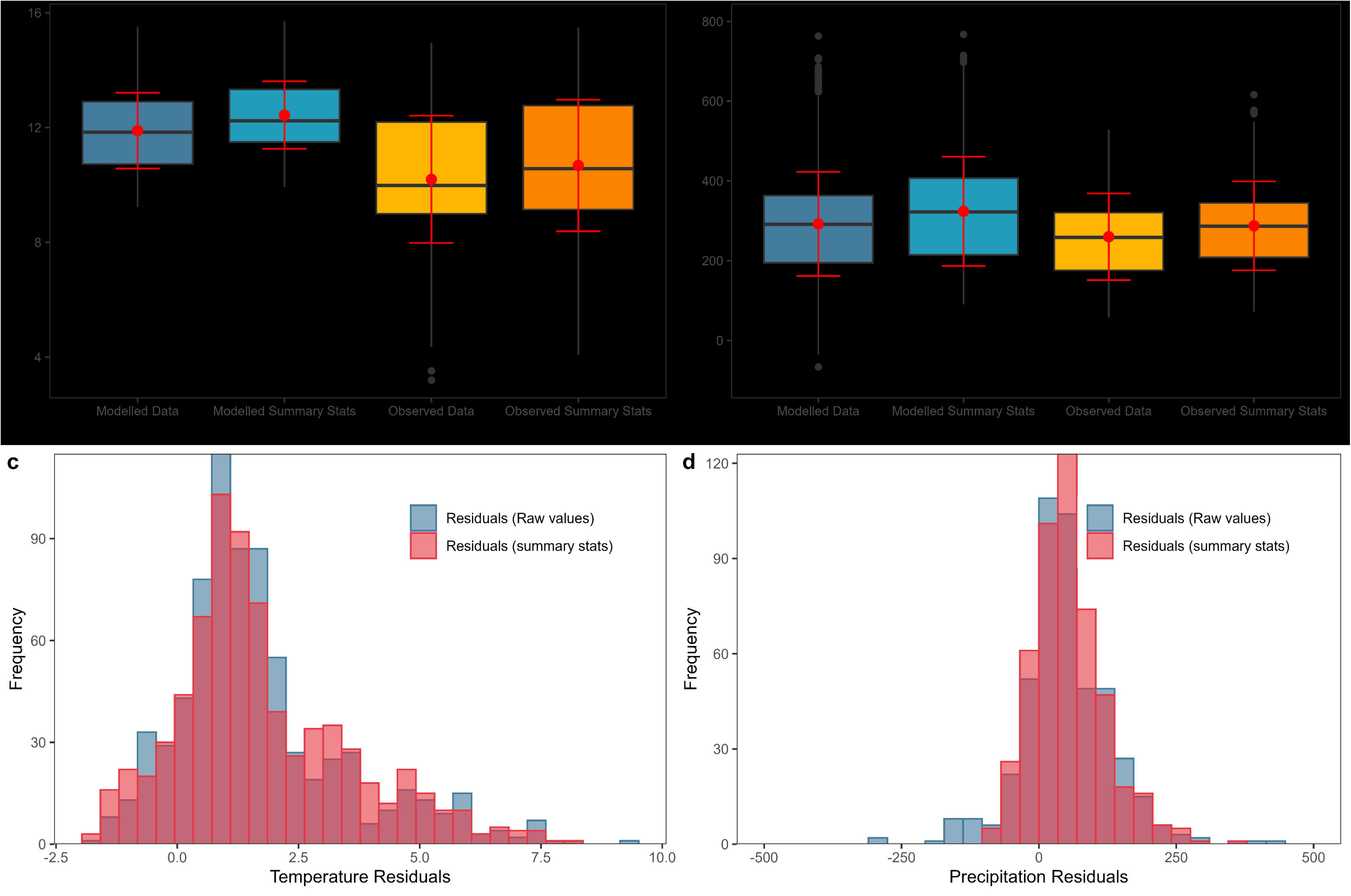
Results of model validation approach, comparing expected shortfall results when calculated from modelled and observed raw data values and summary statistics. a) displays boxplots of calculated expected shortfall values for estuaries in the Pacific Northwest (n = 749) for both modelled and observed temperature data, and b) displays boxplots that contain expected shortfall values for both modelled and observed precipitation data. For both a) and b), boxes represent the median and interquartile range values, while red points are given by mean and red error bars are given by the standard deviation. Black points denote outlier values as defined by the smallest and largest values further than 1.5 times the inter-quartile range from the mean (Wickham et al., 2026). c) plots the residuals of expected shortfall values calculated for temperature data, and d) plots the residuals of expected shortfall values calculated for precipitation data. Plots b and d exclude 18 outlier estuaries for ease of interpretation, for full plots see S3. Modelled data refers to data from the CHELSA downscaled climate models from 1981 to 2020, which exists at a 0.01 degree resolution. Observed data refers to climate data from the NOAA’s global historical climate dataset between 1908 and 2020, which exists at a 0.5 degree resolution.

For precipitation, there was little difference in the calculated expected shortfall between the observed and modelled data, as the means of each calculation method were within one standard deviation of each other (Figure 4b). A bootstrapped Kruskal-Wallis test indicated no significant difference in the results calculated between all 4 datasets (mean Kruskal-Wallis value (df = 3, N = 4) = 3.350953, mean p value = 0.4165543). However, there were 18 estuaries in the Southwest British Columbia region where observed precipitation values were much higher than modelled averages, resulting in expected shortfall values an order of magnitude higher than when predicted using modelled data (2000-8000mm of precipitation, compared to the range of -100-800mm seen in Figure 4b). This indicates a potential underprediction in expected shortfall for those estuaries (see S3 for more information), and resulted in a large difference between the means of the residuals (mean raw value residuals = -142.41 ± 1098.55, mean summary statistic residuals = 54.96 ± 65.13; Figure 4d). Overall, there was a similar overlap in values between the extremes calculated using the raw data and the extremes calculated using the summary statistics for both variables, indicating that the use of summary statistics was appropriate for the analysis here conducted.

### Changes in mean

Across all seven regions, during the mid-century period of 2041-2070, estuaries are projected to experience an increase in mean temperature, with the average estuary experiencing an increase of 22 percent and a median increase of 19 percent. This indicates that the region’s average could increase from around 10□ to around 12□. Consistent with the predictions for SSP scenarios, the results for SSP 1 show a smaller increase in mean temperature when compared to SSP 5. The region with the largest projected increase in mean temperature is North Central BC, which is the region furthest north within the study area, and also contains the most estuaries (n = 207). This region also contains the individual estuaries that are projected to experience the largest increase in mean temperature (including the Hans Point estuary, Figure 6a). In contrast, the region that will experience the least amount of change is Central California, which is the area furthest south within the study area, and the region with the fewest estuaries (n = 31). The difference between most and least amount of change is reflective of the general latitudinal gradient present, where regions further north will experience a stronger increase in mean temperature. On average, the range between maximum and minimum percent change in temperature for estuaries in the Pacific Northwest is 34.4 percentage points.

Trends in mean precipitation projections across estuaries in the Pacific Northwest are not as consistent as temperature projections. On average, estuaries could experience an increase in precipitation of 10.7 percent, and a median increase of 10.1 percent. This translates to an increase from 173 mm/month to 190 mm/month. While the majority of estuaries in the study area will likely experience an increase in mean precipitation under both SSP scenarios, this trend does not hold true for all the estuaries in the study area, as there are some significant exceptions in the Salish Sea and Central California regions where some estuaries may experience no change or a decrease in mean precipitation. There is little difference between both SSP scenarios when it comes to increases in monthly precipitation, as the region will likely only experience a small to moderate increase in monthly precipitation under SSP 5 when compared to SSP 1. Haida Gwaii, the region with the highest increase in mean precipitation, is the only region where all estuaries are expected to increase in monthly precipitation. Every other region has some estuaries that may experience no change or a decrease in precipitation; in particular, many estuaries in the Salish Sea, WA, will likely experience a strong decrease in precipitation. In Central California, while some estuaries may experience little change or a small increase in monthly precipitation, under SSP 5, some estuaries may experience the largest increases in precipitation out of all the estuaries in the study area. On average, the difference between the maximum and minimum mean percent change in precipitation for estuaries in the Pacific Northwest is 13.3 percentage points.

### Changes in variability

While temperature is primarily expected to increase for estuaries in this region (Figure 6), variability in temperature (here given by the standard deviation) displays a different trend. Temperature variability is on average not expected to change much from current conditions, with all estuaries having an average increase in variability of 4.5 percent and a median increase of 3.4 percent. This is synonymous with an increase from a standard deviation of 1.8□ to 1.9□. However, under SSP 5-RCP 8.5, there is a more pronounced increase in temperature variability in estuaries. For estuaries in the northern regions of the study area (Northern BC and Haida Gwaii), temperature variability is centered close to 0, and these estuaries may not experience much change. However, estuaries throughout the rest of the study area may experience a slight increase in temperature variability, with the southernmost estuaries in Central California experiencing the strongest increase. This trend is the opposite of changes in mean temperature, where southern regions are predicted to become more variable, while northern regions may become more stable. On average, the difference between the maximum and minimum change in temperature variability in estuaries in the Pacific Northwest is 25.1 percentage points.

The mean percent change in precipitation variability is -11 percent, and the median is - 11.1 percent, indicating a general decrease in precipitation variability from a standard deviation of 79 mm/month to 71 mm/month. Interestingly, this trend is consistent across both SSP scenarios included in this analysis, as there is little variation between the results in either SSP scenario. On average, the difference between the maximum and minimum amount of change an estuary can experience in precipitation variability is 18.3 percentage points; when compared with the mean and median change value, this indicates that many estuaries can experience a range of effects, and the directionality of that effect may vary.

### Changes in extremes

The magnitude of extreme high temperatures, calculated as the expected shortfall of temperatures in the 99th percentile of mean monthly temperatures, are primarily expected to increase in estuaries across the Pacific Northwest. On average, the magnitude of an extreme temperature event may increase by 10.6 percent, or by a median amount of 9.2 percent. This means that if the average heatwave across the region resulted in a high of 25□, this would mean the average heatwave would increase to 28□. This is consistent with trends in increases in mean monthly temperatures. The northernmost region in the study area, north central BC, could experience the least change under both SSP scenarios compared to other regions, but still displays a relatively wide range of effects for the estuaries across its region, consistent with the other regions in the study area. Estuaries in central CA (at the southernmost range of the study area) will likely experience the largest increase in extreme temperatures, and this region contains some of the most extreme outliers.

When it comes to changes in magnitude of extreme precipitation, estuaries in the Pacific Northwest will generally experience a decrease, as the mean percent change value is -4.7 percent, and the median is -4.9 percent. If the average extreme rainfall across the region is around 328 mm/month, this would mean the average extreme rainfall event would decrease to 312 mm/month. These trends closely follow changes in precipitation variability (Figure 6).

Under SSP 1, most estuaries may experience a decrease in extreme precipitation, with estuaries in Central California experiencing the strongest decrease. Northwest Vancouver Island presents a notable exception, where estuaries may experience either an increase or a decrease in extreme precipitation. Under SSP 5, this region may experience an increase in extreme precipitation on average, while the remaining regions may experience a decrease (with some notable strong exceptions in Haida Gwaii). Interestingly, under SSP 5, estuaries may generally experience less of a decrease in extreme precipitation than they would under SSP 1.

## Discussion

### Describing complex climate futures for conservation can be done using accessible approaches

Our research sought to describe changes in the mean, variability, and magnitude of extremes in temperature and precipitation for ecological management, using a case study of 749 estuaries in the Pacific Northwest. Our approach for describing these effects as a percent change from a historical baseline is straightforward and makes use of existing open access climate projection datasets. Our approach used a popular open-source software (RStudio); however, these indicators can be calculated using other software (eg. MS Excel) and offer relatively stable estimates of change. Importantly, these can be easily communicated and interpreted by conservation managers, allowing them to create more robust management plans.

Using global downscaled open source datasets such as CHELSA also allows managers in other regions who may not have access to local-scale climate projections to still apply this approach. Local scale climate projections are key to this approach, as temperature and precipitation depend much more on geographical patterns that can affect local weather systems. For example, our results present a general trend of increase in both mean temperature and mean precipitation for estuaries in the Pacific Northwest, which is consistent with global-scale climate projections (Intergovernmental Panel On Climate Change (IPCC), 2023), but estuaries such as the inner reaches of the Columbia River, which may experience an increase in mean temperature (between 9.1 and 36.9 percent) but a potential decrease in mean precipitation (between -12.2 and 9.9 percent) don’t follow this global trend. When examining variability and magnitude of extreme events, this becomes more evident, as global projections do not match local trends. For example, precipitation variability is expected to increase globally (Tahroudi, 2025; Pendergrass et al., 2017), but our results suggest an overall decrease in precipitation variability in this region (Figure 6d). At the regional scale, using data with a 0.01 degree resolution (approximately 1 square kilometre) is appropriate; however, for a smaller region or a more local scale analysis, finer scale projection data might be more appropriate. The approach described here shows promise at the regional scale and may be able to be adapted to smaller scales, but more study is needed in this area.

With increasing availability of downscaled climate projection models, there is still a lack of local-level projections of extreme events due to the complexities inherent in predicting these, as they are rare by definition. In our methods, we also outlined a novel method of projecting the magnitude of extreme events using extreme value theory, which relies only on open source projections of mean and variability. Given the limited availability of climate projection data at a fine temporal resolution (eg. monthly or yearly data), our approach of using summary statistics potentially overpredicts the magnitude of extreme events in temperature (Figure 4a, 4c, S4), resulting in a significant difference in the expected shortfall when calculated using modelled summary statistics compared to using observed individual monthly values (S4). However, by overpredicting rather than underpredicting, the model errs on the side of caution; use of the worst-case-scenario for conservation planning can result in better conservation outcomes (eg. Maes et al., 2022; O’Hanley et al., 2007). This method, alongside estimates of changes in mean and variability, can provide a more comprehensive overview of the effects of climate change on an ecosystem, while working to counter some of the limitations of climate projection data.

A well recognized shortcoming of climate projections are the inherent uncertainties in climate projection models (Frigg et al., 2015). Rather than attempt to reduce uncertainty, our approach leans into examining all possible futures as a form of improving preparedness for future possibilities. Examining a wider range of futures by providing 10 predicted values (5 global climate models each for 2 SSP scenarios) for each variable rather than a single mean value allows us to detect discrepancies in predictions, where different climate models may predict change in opposing directions, as seen in the changes in temperature variability described for the Hiellen River in Figure 7a. If temperature variability is projected to change by anywhere between -14.8 percent and 12.1 percent, it can be tempting to reduce this range to a mean value. However, planning that includes or acknowledges uncertainty can be more robust (Blankespoor et al., 2023), and acknowledges that decision makers may have different levels of risk aversion and may seek to tailor scientific outputs to different decision making contexts (Douglas & Wildavsky, 1982). For example, more risk-averse managers may be more concerned with the possibility of increased temperature variability, which is linked to changes in plant growth timings (Thornton et al., 2014) and impacts ecological interactions (Caparros-Santiago et al., 2021; Visser, 2016). Thus, managers may elect to incorporate restoration activities that strengthen plant communities in the Hiellen River estuary (e.g., management of introduced Sitka Black-tailed Deer, *Odocoileus hemionus sitkensis,* that graze on native plants, *Secretariat of Haida Nation, n.d.)* to ensure a greater diversity of native plant species can still fill their required ecological niche.

This project makes use of some of the more extreme projections for climate (SSP 5-8.5) that may seem implausible, as they focus on energy economics and climate forcing, and rely heavily on inputs such as global coal use (Pielke Jr et al., 2022; Ritchie & Dowlatabadi, 2017). Using these scenarios, our study potentially overpredicts the magnitude of extreme events for some estuaries (Figure 4). Despite this, because conservation is concerned more with effects than exact predictions, the outputs of these models are still useful for climate adaptation strategies (Clark-Wolf et al., 2025; Littell et al., 2011; Peterson et al., 2014). This approach can provide more risk-averse conservation planners with the security of knowing they can over-prepare for the worst case scenario. This knowledge can strengthen post-event recovery plans, improving adaptive capacity (Field et al., 2012). By examining all predicted outcomes, it allows conservation planners to consider which changes would likely have a more detrimental effect on their ecosystem, and plan accordingly.

### Examining changes in mean, variability, and magnitude of extremes together highlight diverse impacts to Pacific Northwest estuaries

The results of this study highlight the importance of understanding changes in mean, variability, and magnitude of extremes of climate variables together. Based on a case study of estuaries in the Pacific Northwest, the results highlight diverse impacts that will paint a more complete picture of the local effects of climate change. While this study only focused on one ecosystem type, changes in mean, variability, and extremes have been seen to be ubiquitous across ecosystems (Field et al., 2012; Rodgers et al., 2021), making this framework of understanding climate effects potentially useful across various ecosystems.

Because the calculation of standard deviation is dependent on the mean, changes in both of these statistics often occur in the same direction. However, this may not be the case in ecological systems, as exemplified here by the case study of estuaries. Despite an overall increase in mean temperature (Figure 5b), temperature variability may decrease for many of the estuaries present in the study area (Figure 6b). This could indicate a shift toward a more stable, less variable climate, but can also indicate more mild winters, and warmer spring and fall seasons, as the temperature range of a region narrows, as described by a decrease in standard deviation (variability). Similarly, the projected decrease in precipitation variability alongside an expected increase in mean precipitation could indicate a decrease in precipitation during the spring and fall, which are the periods when precipitation is most variable in the region. These seasonal precipitation patterns are important for nutrient flow, tidal connection, and aquatic species such as anadromous salmon (*Oncorhynchus spp.*), which are cultural keystone species in the Pacific Northwest (L. A. Jones et al., 2020; Poirier et al., 2012). For example, for large estuaries such as Humboldt Bay, while mean precipitation is predicted to increase (between 16.9 and -0.5 percent), the predicted decrease in precipitation variability (between -2.03 and -23.3 percent) could bring challenges for local salmon and other seasonal precipitation-dependent species (Siegel & Crozier, 2019; Wittig & Déry, 2026). As inherently variable ecosystems, estuarine conditions are tied to seasonal patterns in temperature and precipitation, which drive salinity, streamflow, primary productivity, plant community cycles, and migratory patterns for many species (Chilton et al., 2021; Snow et al., 2000). While many estuarine species are adapted for variability, changes in seasonal patterns due to climate change occur at a rate that is unsustainable for adaptation (Cloern et al., 2016), making it essential to consider changes in variability alongside changes in mean in conservation planning.

**Figure 5:**
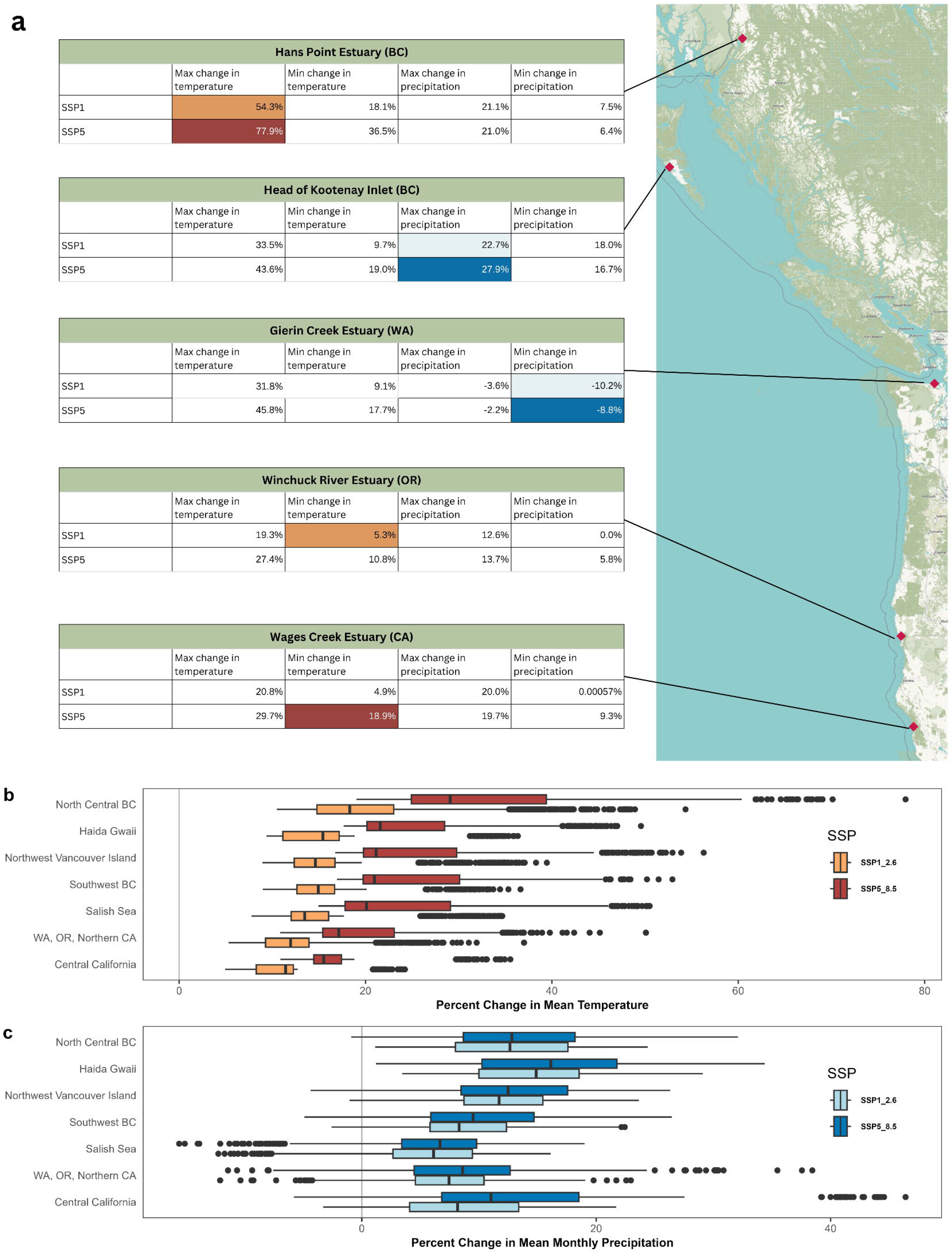
Percent change in mean monthly temperature and monthly precipitation from historical baseline (1980-2011) to predicted futures in the mid-twenty-first century (2041-2070) across estuaries in the Pacific Northwest (n = 749). (a) maps of estuaries that will experience the most and least change in mean, where colored boxes indicate the value is the highest or lowest in its category. (b) shows the mean and inter-quartile range of percent change in mean temperature at two SSP scenarios at seven sub-regions of the Pacific Northwest, and (c) shows the mean and inter-quartile range of percent change in mean monthly precipitation. Black points denote outlier values as defined by the smallest and largest values further than 1.5 times the inter-quartile range from the mean (Wickham et al., 2026).

Trends in projected changes in mean and magnitude of extreme events may differ in magnitude, making these trends essential to understand together given the ecological effects of extreme events in temperature and precipitation. In line with global climate projections, our results outline a parallel increase in the mean and magnitude of extreme events in temperature (Figure 5b & Figure 7b). Examining trends in means and extremes together reveals interesting trends— for example, while Central California is projected to experience the smallest increase in mean temperature compared to other studied regions (between 4.92 percent and 35.53 percent; Figure 5b), it is projected to experience the largest increase in magnitude of extreme temperatures (between -0.38 percent and 44.38 percent; Figure 7b). These extreme events in temperature (*e.g.*, heat waves) are ecologically significant, as they can create ecosystem-wide impacts such as species mortality, behavioural changes, exposure to disease, and changes in community assemblages (Anadón et al., 2024; Bateman et al., 2020; Field et al., 2012; Pérez-Romero et al., 2019; Shields et al., 2019). While the specific effects of these events on individual estuaries will depend on their unique hydrological and elevation profiles (Winter, 2000), examining these trends together can help individual managers, practitioners, and knowledge holders to understand and plan for change, using strategies to mitigate these effects. For example, increased tillage or reintroduction of cultural fire practices can improve biodiversity and community resilience to hotter, dryer periods (Xiong et al., 2023).

Trends in projected change in mean and magnitude of extreme events may also differ in direction. Interestingly, our results showed a projected decrease in the magnitude of extreme precipitation (Figure 7c), despite a projected increase in mean precipitation (Figure 5c). This projected change may be seen positively, as extreme precipitation events (*e.g.*, extreme rainfall, storm surges, or drought) can lead to changes in salinity, changes in sediment transportation and accretion, and changes in nutrient flow (Robins et al., 2016; Towler et al., 2010). However, many estuarine ecosystems are adapted to rely on high rainfall events in the fall and winter (Dettinger, 2011; Sobral & Déry, 2023), so a significant decrease in extreme rainfall may result in negative effects on estuarine organisms such as anadromous salmon (Wittig & Déry, 2026). Understanding how these changes in magnitude of extreme events may affect ecosystems such as estuaries differently is an important strategy to guide activities such as restoration of habitat for improved water retention and streamflow (Chilton et al., 2021).

Trends in variability can also provide insight into the frequency of extreme events in both temperature and precipitation. While our data prevented us from explicitly examining frequency of extreme events (daily data would have provided better information, as extreme events don’t frequently occur on the monthly scale), changes in the standard deviation of a climate variable also indicates changes in the frequency of occurrence of events that today would be considered extreme. As this region is projected to primarily experience increased temperature variability (Figure 6b), this could indicate increased frequency of extreme events in temperature. Similarly, decreased precipitation variability (Figure 6c) could indicate decreased frequency of extreme events in precipitation. However, further research is needed to better quantify the frequency of extreme events in estuaries of the Pacific Northwest and across ecosystems.

**Figure 6:**
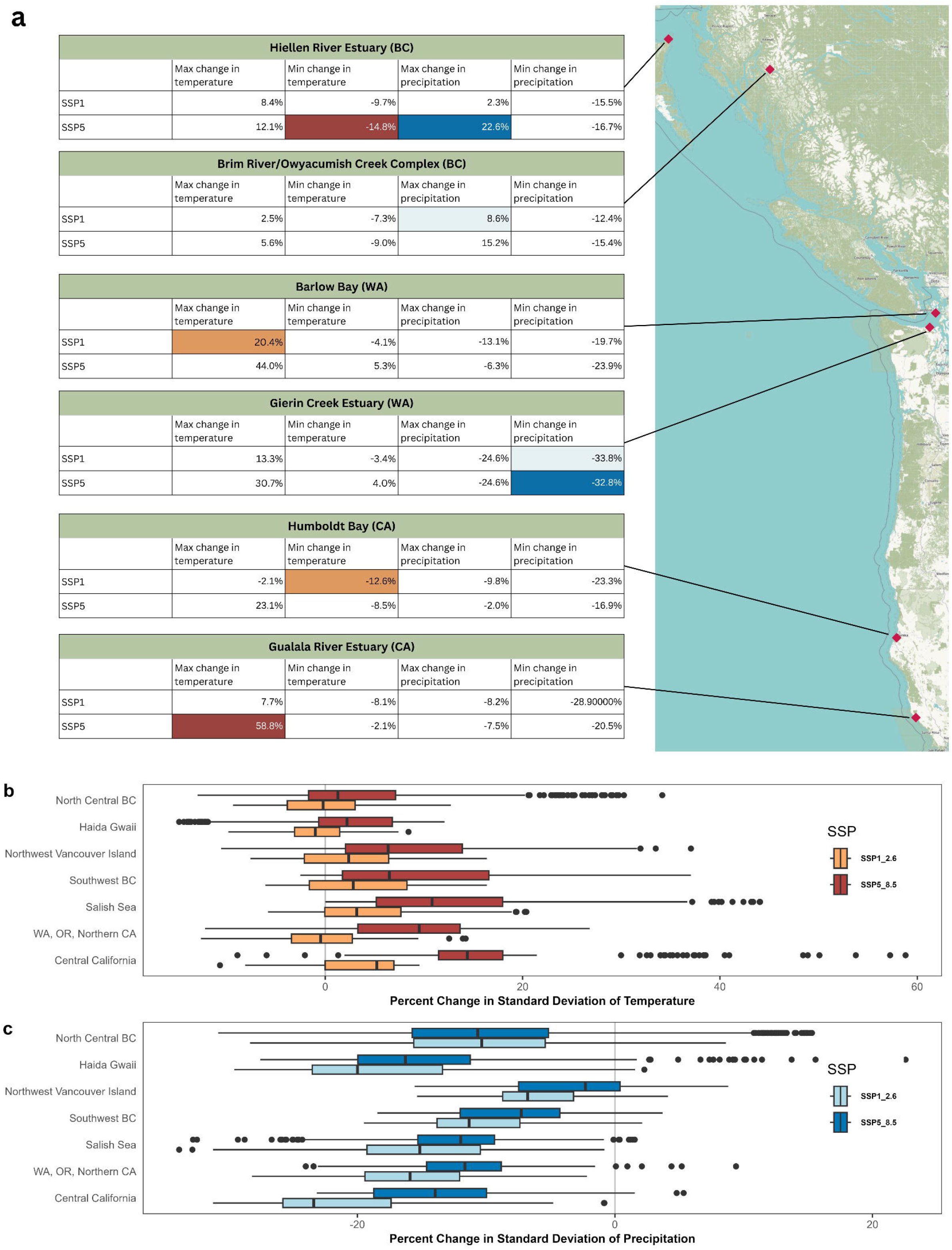
Percent change in standard deviation of mean monthly temperature and standard deviation of monthly precipitation from historical baseline (1980-2011) to predicted futures in the mid-twenty-first century (2041-2070) across estuaries in the Pacific Northwest (n = 749). (a) maps estuaries who will experience the most and least change in standard deviation, where colored boxes indicate the value is the highest or lowest in its category. (b) shows the mean and inter-quartile range of percent change in standard deviation of temperature at 2 SSP scenarios at seven sub-regions of the Pacific Northwest, and (c) shows the mean and inter-quartile range of percent change in standard deviation of monthly precipitation. Black points denote outlier values as defined by the smallest and largest values further than 1.5 times the inter-quartile range from the mean (Wickham et al., 2026).

### Understanding different dimensions of climate change is essential to improving conservation planning

Many have acknowledged that climate change is an important part of modern day conservation planning (Halperin et al., 2024; Hannah et al., 2002b; Marcus et al., 2026). Some approaches to climate-informed conservation planning include mapping species distribution ranges under climate change, mapping climate refugia as areas for protection, improving connectivity between habitats, and regional modelling and resource coordination (Hannah et al., 2002a; Hansen et al., 2010; K. R. Jones et al., 2016; Liberati et al., 2016). However, many of these strategies don’t examine trends in mean, variability, and magnitude of extremes simultaneously, which may lead to underestimating climate effects and reactionary rather than proactive planning. By studying these trends together, conservation planners may have a better idea of what conservation strategies are most appropriate for each region, and better prioritize resources for more effective biodiversity conservation.

An important conservation strategy is ecological restoration, which includes habitat remediation and enhancement, especially in degraded regions. A potential application of this work is to identify regions that are at risk of increased climate impacts, and planning restoration work that is in line with projected climate effects. For example, the Olympic Mountains rain shadow (Pater et al., n.d.) has a strong effect on many estuaries in the Salish Sea and Southern British Columbia regions. This rain shadow is reflected in the results here, as some estuaries in this rain shadow— such as Gierin Creek in Washington—are predicted to experience a decrease in mean precipitation (between -2.2 and -10.1 percent), despite the average for the region predicting an increase (Figure 5a). This estuary is also predicted to experience a decrease in precipitation variability (between -24.5 and -33.8 percent, Figure 6a) and a decrease in magnitude of extreme precipitation (between -19.5 and -27.8 percent, Figure 7a). Gierin Creek has been identified as an area for conservation action—not because of projected climate actions, but due to impacts from agricultural runoff, installation of a tidal gate that converted saltmarsh to freshwater wetland, and the lack of large woody debris (LWD; Elwha-Dungeness Planning Unit, 2005). Recommended actions for restoration include removal of the tidal gate and addition of LWD to add nutrients to the soil (Elwha-Dungeness Planning Unit, 2005), both of which are also strategies that could help the estuary adapt to a potentially hotter and dryer future by improving moisture retention, creating shade for invertebrates and other aquatic species, and improving plant community health. Overall, better knowledge of climate impacts can help tailor restoration projects toward improved resiliency and support practitioners in prioritizing resources for existing projects. Examples such as improvement or restoration of hydrological flow (Riley et al., 2018), creation of habitat for vulnerable species (Camp et al., 2015; Gaines et al., 2024), or vulnerable species harvesting limits (Camp et al., 2015) are good examples of restoration actions that take into account climate impacts. However, extreme climate event-resilient restoration remains an understudied field, and restoration actions will likely vary considerably across contexts.

**Figure 7:**
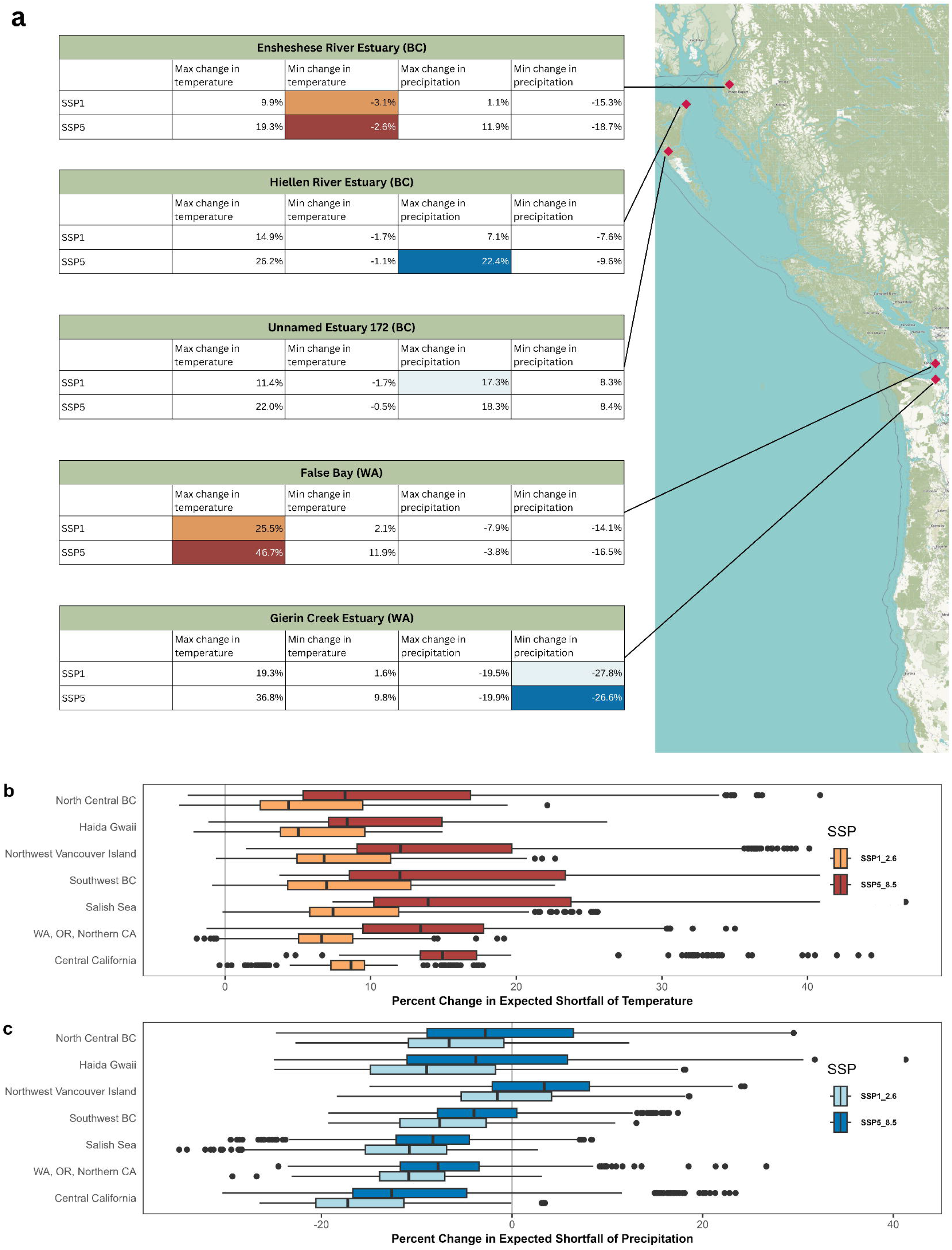
Percent change in expected shortfall at the 99th percentile of monthly temperature and monthly precipitation from historical baseline (1980-2011) to predicted futures in the mid-twenty-first century (2041-2070) across estuaries in the Pacific Northwest (n = 749). (a) maps estuaries who will experience the most and least change in expected shortfall, where colored boxes indicate the value is the highest or lowest in its category. (b) shows the mean and inter-quartile range of percent change in expected shortfall of temperature at 2 SSP scenarios at seven sub-regions of the Pacific Northwest, and (c) shows the mean and inter-quartile range of percent change in expected shortfall of monthly precipitation. Black points denote outlier values as defined by the smallest and largest values further than 1.5 times the inter-quartile range from the mean (Wickham et al., 2026).

Another focus of many regional conservation planners is land acquisition, much of which is then converted to nature preserves, protected areas, or other effective conservation measures. While many factors go into consideration for land acquisitions, including species targets and human activity (K. R. Jones et al., 2016; Robb, 2025), knowing how an area may be impacted by climate change is valuable to ensure conservation goals are being met into the future. For example, a conservation planner in British Columbia may be concerned with the impacts of heatwaves on salmon populations, and may wish to conserve areas that may not experience heatwaves as strongly, and thus can act as refugia for salmon. Our results predict that Younghpan Creek, located in the Holberg Inlet on Vancouver Island, may not experience as strong an increase in magnitude of extreme temperatures compared to other estuaries (between 0.01 and 20 percent). Currently, this estuary is not protected under any conservation measures, but has been identified as valuable habitat for both migratory birds (Pacific Birds Habitat Joint Venture Technical Team, 2019) and salmon species (Brown et al., 1979), including Coho (*Oncorhynchus kisutch*) and Chum (*Oncorhynchus keta*). Additionally, it has been identified as a region that experiences increased pressures from human activity, including forestry and recreational and commercial fishing (Robb, 2025). By combining climate data with other conservation related goals, regional conservation planners may be able to better identify and prioritize conservation of climate refugia.

By examining trends in mean, variability, and magnitude of extreme events of important climate variables, conservation planners can obtain a robust yet non-deterministic estimate of how their sites may change in the future. Especially at the regional scale, the method here outlined can help large conservation organizations (eg. The Nature Conservancy, Ducks Unlimited, Pacific Birds Habitat Joint Venture) to prioritize resources and capacity for all types of conservation action. This method stands out as key for conservation planning that considers not only mean changes but planning for the variable future that climate change promises.

## Conclusion

Climate-integrated conservation planning is a growing area of research, and many researchers and conservation practitioners alike have put forth valuable frameworks for integrating climate projections into action. The work here presented expands on these ideas by providing a straightforward method of examining changes in mean, variability, and extremes in climate projections, all of which are ecologically significant and may affect ecosystems in different ways. This framework fills the gap between climate knowledge and conservation planning, breaking down climate projection information in a way that can better inform conservation practitioners toward a climate resilient future. For conservation planning that goes beyond the local level and hopes to protect flyways, migratory pathways, and regional connectivity, accessibility of climate information in a straightforward, usable form for practitioners will become increasingly important.

This research presents several areas for further research. First, by pairing this information with other biotic metrics such as diversity and species composition; abiotic metrics such as hydrological regimes, elevation, and aspect; or metrics of cultural or economic importance, these data can provide regional managers with the information to effectively prioritize estuaries for conservation action. Additionally, the methods here presented show promise, and could potentially be adapted to other ecosystem contexts and climate variables. Across ecosystems, understanding how our lands may change in the future can help to ensure effective conservation action for decades to come.

## Acknowledgements

This work was funded by a Mitacs Accelerate Research Fellowship. We thank the Nippon Foundation Ocean Nexus for their support. We would also like to acknowledge that this research was conducted on the traditional territories of the lək□□əŋən and W□SÁNEĆ peoples, and this research covers an area stewarded by over 330 First Nations and Tribes, who have cared for this land since time immemorial.

## Author Contributions

RM: conceptualization, analysis, writing — original draft, writing — review and editing. NS: conceptualization, project administration, writing — review and editing. GS: conceptualization, project administration, writing — review and editing.

## Conflict of Interest

The authors declare no conflicts of interest.

## Data Availability

All data other than BC estuary shapefiles are open source and publicly accessible. Estuary shapefiles for WA, OR, and CA can be found at https://www.pacificfishhabitat.org/data/estuary-extents. CHELSA climate data, including historical and projected mean and standard deviation of temperature and precipitation, can be found at https://www.chelsa-climate.org/. NOAA global gridded monthly land temperature data can be found at https://psl.noaa.gov/rest/data.ghcncams.html. NOAA University of Delaware global monthly terrestrial precipitation data can be found at https://psl.noaa.gov/data/gridded/data.UDel_AirT_Precip.html. Results data as generated by this project are available via GitHub at https://github.com/rekhamarcus/PNW_Estuaries.

## Code Availability

The code used to generate the dataset, analysis, and figures can be found in the GitHub repository at https://github.com/rekhamarcus/PNW_Estuaries.

## Supplementary Materials

S1: Projected change in expected shortfall at 95th percentile

S2: Model ensemble results

S3: Mapped estuaries where modelled precipitation projections don’t match observations

## Notes

https://github.com/rekhamarcus/PNW_Estuaries

### Competing Interest Statement

The authors have declared no competing interest.

