## Supplemental Figure 1 for "Characterizing multiple dimensions of climate hazards for conservation planning: a case study of estuaries in the Pacific Northwest"

### S1: Projected change in expected shortfall at 95th percentile

**a**

| Ensheshese River Estuary (BC) | | | | |
|---|---|---|---|---|
| | Max change in temperature | Min change in temperature | Max change in precipitation | Min change in precipitation |
| SSP1 | 12.2% | -2.1% | 2.8% | -13.4% |
| SSP5 | 21.9% | -0.6% | 12.8% | -16.8% |

| Oeanda River Estuary (BC) | | | | |
|---|---|---|---|---|
| | Max change in temperature | Min change in temperature | Max change in precipitation | Min change in precipitation |
| SSP1 | 16.5% | -0.2% | 7.4% | -6.7% |
| SSP5 | 25.5% | 1.4% | 22.4% | -8.5% |

| Unnamed Estuary 172 (BC) | | | | |
|---|---|---|---|---|
| | Max change in temperature | Min change in temperature | Max change in precipitation | Min change in precipitation |
| SSP1 | 13.5% | -0.5% | 17.5% | 8.2% |
| SSP5 | 24.1% | 1.6% | 18.3% | 8.4% |

| False Bay (WA) | | | | |
|---|---|---|---|---|
| | Max change in temperature | Min change in temperature | Max change in precipitation | Min change in precipitation |
| SSP1 | 27.5% | 3.1% | -6.9% | -12.9% |
| SSP5 | 47.1% | 12.9% | -2.9% | -15.2% |

| Gierin Creek Estuary (WA) | | | | |
|---|---|---|---|---|
| | Max change in temperature | Min change in temperature | Max change in precipitation | Min change in precipitation |
| SSP1 | 20.6% | 2.4% | -18.5% | -26.6% |
| SSP5 | 37.7% | 10.6% | -18.7% | -25.5% |

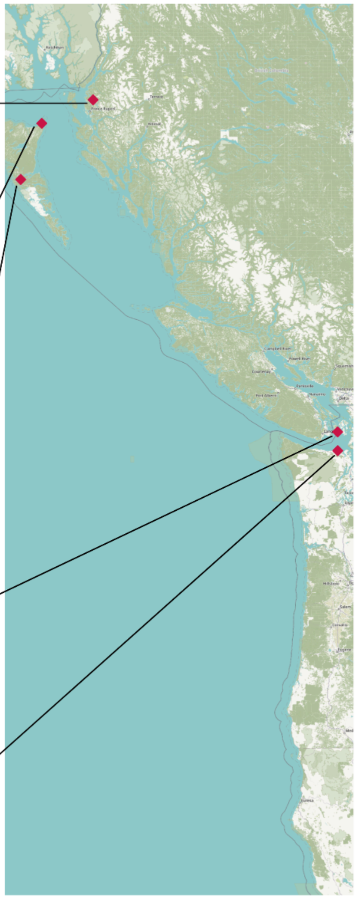

**b**

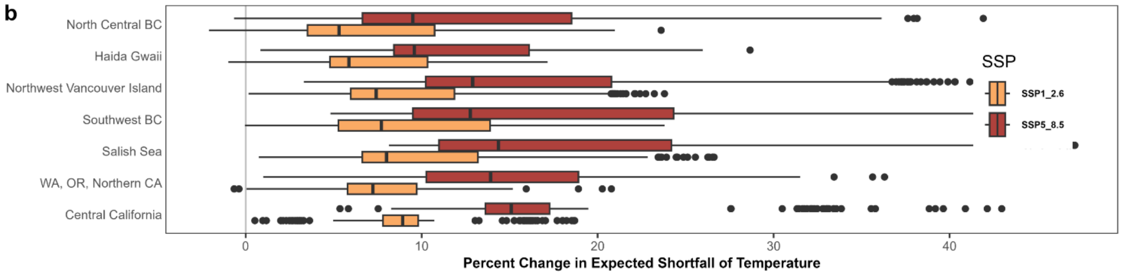


**c**

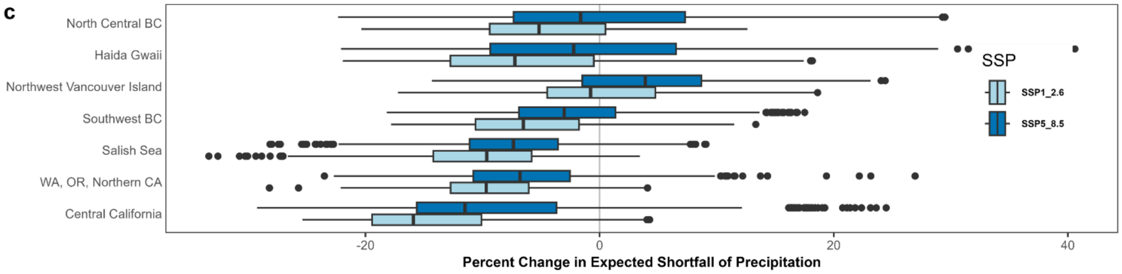

*Figure S1: Percent change in expected shortfall at the 95th percentile of monthly temperature and monthly precipitation from historical baseline (1980-2011) to predicted futures in the mid-twenty-first century (2041-2070) across estuaries in the Pacific Northwest (n = 749). (a) maps estuaries who will experience the most and least change in expected shortfall, where colored boxes indicate the value is the highest or lowest in its category. (b) shows the mean and inter-quartile range of percent change in expected shortfall of temperature at 2 SSP scenarios at seven sub-regions of the Pacific Northwest, and (c) shows the mean and inter-quartile range of percent change in expected shortfall of monthly precipitation. Black points denote outlier values as defined by the smallest and largest values further than 1.5 times the inter-quartile range from the mean* (Wickham et al., 2026).
