## Supplemental Figure 2 for "Characterizing multiple dimensions of climate hazards for conservation planning: a case study of estuaries in the Pacific Northwest"

S2: Model ensemble results

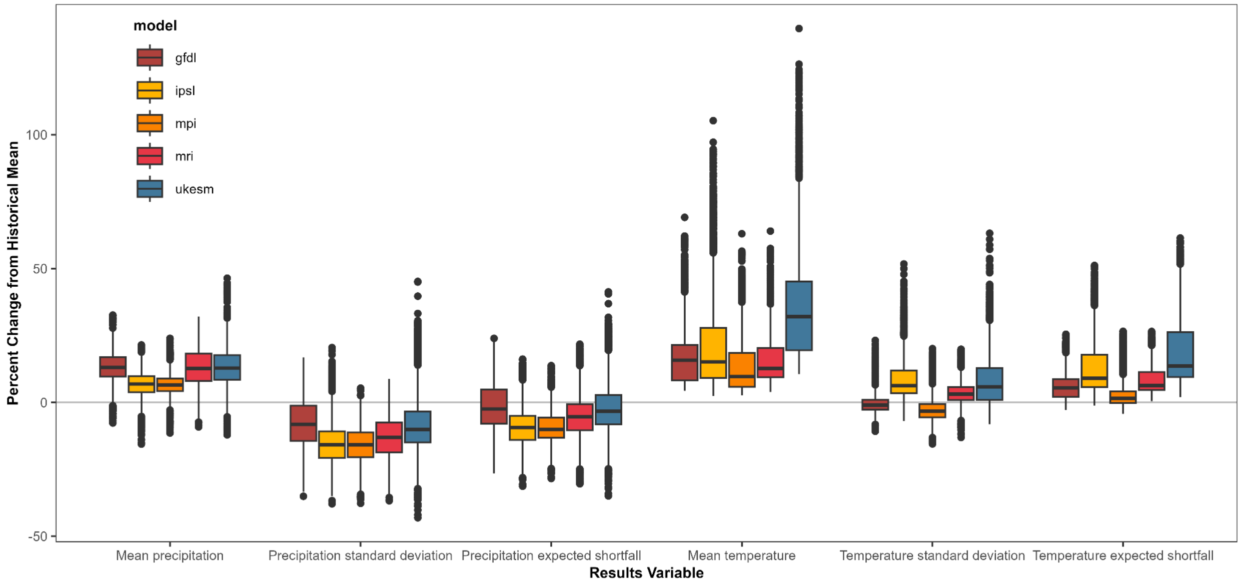


*Figure S2: Plotted results of percent change from historical mean produced from 5 different modelled projection datasets. Black points denote outlier values as defined by the smallest and largest values further than 1.5 times the inter-quartile range from the mean (Wickham et al., 2026).*
