## Supplemental Figure 3 for "Characterizing multiple dimensions of climate hazards for conservation planning: a case study of estuaries in the Pacific Northwest"

<u>S3: Mapped estuaries where modelled precipitation projections don’t match observations</u>

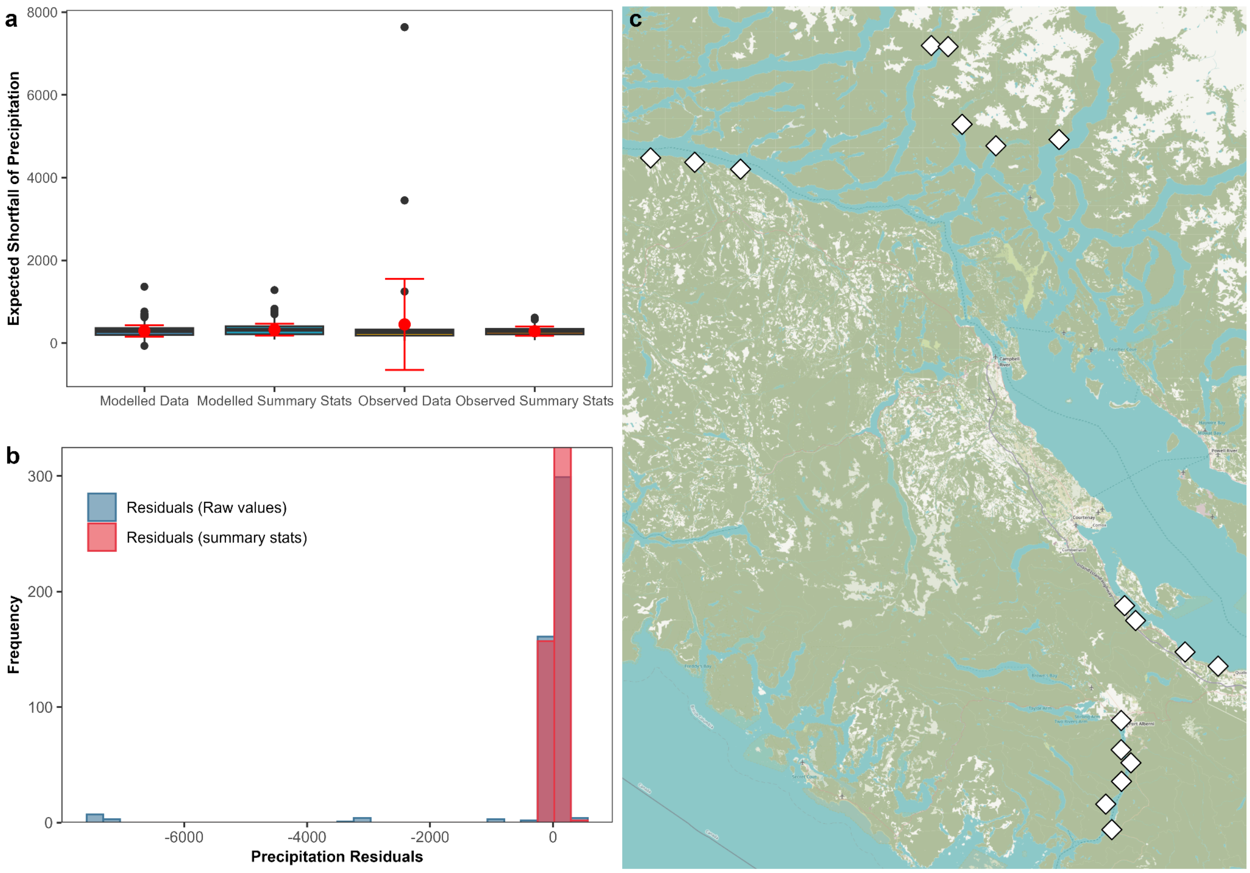


*Figure S3: a) Expected shortfall of precipitation at the 99th percentile as calculated using the 4 datasets described on the x axis; black lines denote the median and inter-quartile range, while the red point denotes the mean and red error bars denote standard deviation. Black points denote outlier values as defined by the smallest and largest values further than 1.5 times the inter-quartile range from the mean (Wickham et al., 2026). b) Residuals of expected shortfall of precipitation calculated using the difference between expected and observed data, using both raw data values and summary statistics. c) mapped estuaries (n = 18, 2.3% of dataset) where modelled historical data (CHELSA) does not match observed historical data (NOAA), resulting in a calculation of expected shortfall that underpredicts the magnitude of extreme precipitation events between 1980 and 2010. Results of expected shortfall for the estuaries mapped resulted in residuals lower than -500mm of precipitation. White diamonds denote estuaries.*
