## Supplemental Figure 4 for "Characterizing multiple dimensions of climate hazards for conservation planning: a case study of estuaries in the Pacific Northwest"

S4: Results of bootstrapped Kruskal-Wallis test for validation data

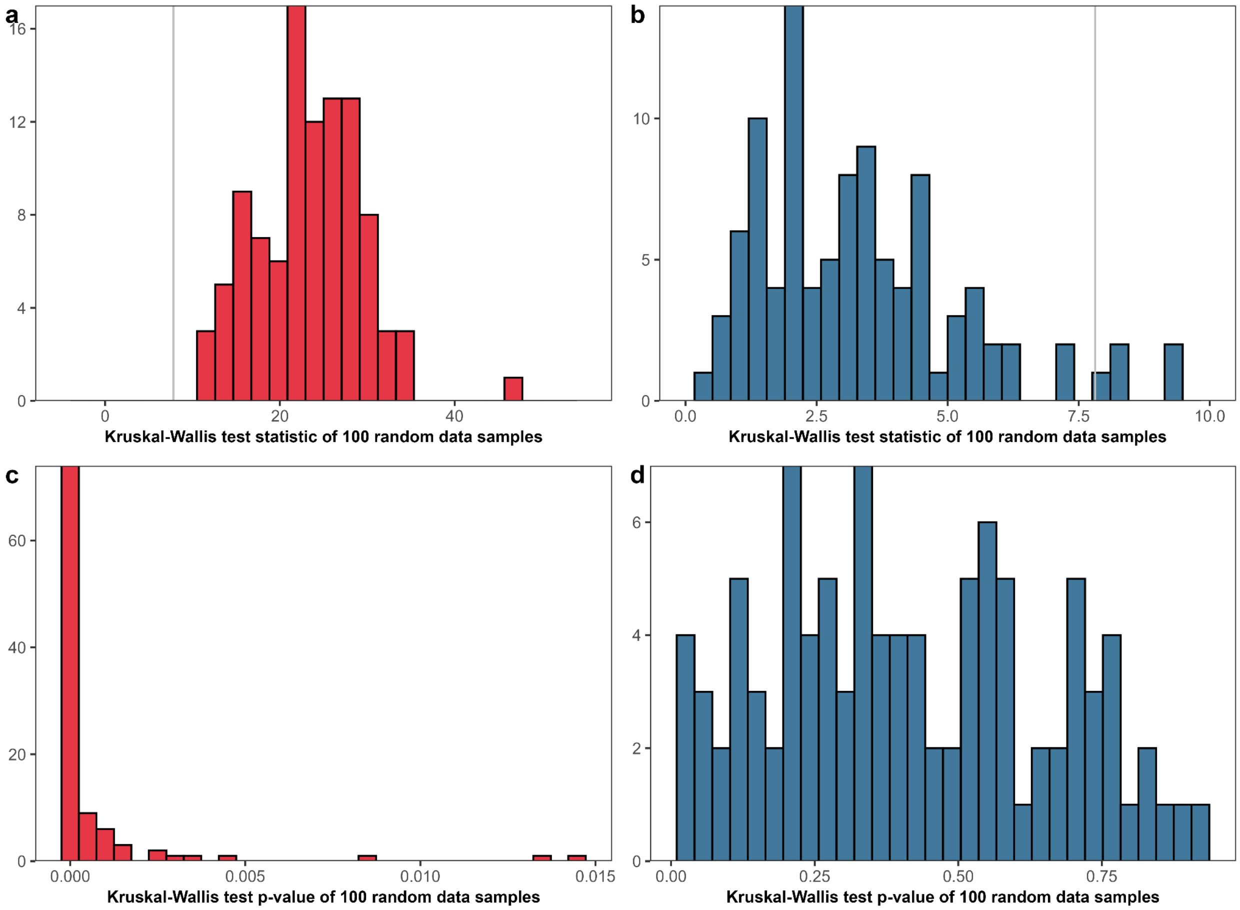


*Figure S4: results of calculating the Kruskal-Wallis test statistic for expected shortfall calculated using 4 different datasets. The test statistics were calculated for a random subset of 30 estuaries, bootstrapped to run 100 times. Panel a) shows the results of the Kruskal-Wallis test statistic for expected shortfall of temperature, where the gray vertical line represents the critical value for a significance level of 5 percent. Panel b) shows the results of the Kruskal-Wallis test statistic for expected shortfall of precipitation, where the gray vertical line represents the critical value for a significance level of 5 percent. Panel c) shows the p-values associated with the calculated Kruskal-Wallis statistics for expected shortfall of temperature, and panel d) shows the p-values associated with the calculated Kruskal-Wallis statistics for expected shortfall of precipitation.*
